# Correlating the structure and activity of *Y. pestis* Ail in a bacterial cell envelope

**DOI:** 10.1101/2020.09.23.310672

**Authors:** J. E. Kent, L. M. Fujimoto, K. Shin, C. Singh, Y. Yao, S. Park, S. J. Opella, G. V. Plano, F. M. Marassi

## Abstract

Understanding microbe-host interactions at the molecular level is a major goal of fundamental biology and therapeutic drug development. Structural biology strives to capture biomolecular structures in action, but the samples are often highly simplified versions of the complex native environment. Here we present an *E. coli* model system that allows us to probe the structure and function of Ail, the major surface protein of the deadly pathogen *Yersinia pestis.* We show that cell surface expression of Ail produces *Y. pestis* virulence phenotypes in *E. coli,* including resistance to human serum, co-sedimentation of human vitronectin and pellicle formation. Moreover, isolated bacterial cell envelopes, encompassing inner and outer membranes, yield high-resolution solid-state nuclear magnetic resonance (NMR) spectra that reflect the structure of Ail and reveal Ail sites that are sensitive to the bacterial membrane environment and involved in the interactions with human serum components. The data capture the structure and function of Ail in a bacterial outer membrane and set the stage for probing its interactions with the complex milieu of immune response proteins present in human serum.

**SIGNIFICANCE:** Ail is a critical virulence factor of *Y. pestis,* and its interactions with human serum are central for promoting the immune resistance of bacteria to the human host defenses. Here we capture the action of Ail in a functional bacterial environment and set the stage for probing its interactions with the complex milieu of immune response proteins present in human serum. The development of an *E. coli* model system of *Y. pestis* for biophysical studies is new and biologically important. Finally, the work extends the range *in-situ* NMR spectroscopy to include models of microbial infection.

## INTRODUCTION

The protein and lipid components of bacterial outer membranes work together to support microbial cell survival in a wide range of host environments and represent key virulence factors. This phenomenon is especially striking in the case of *Yersinia pestis* – the agent responsible for multiple devastating human plague pandemics throughout history. The *Y. pestis* outer membrane protein Adhesion invasion locus (Ail) and lipopolysaccharide (LPS) have co-evolved to enhance microbial resistance to human innate immunity, enabling the bacterium to produce high-level septicemia in its mammalian hosts (1–11). Ail and LPS jointly promote evasion of the host immune defenses and adhesion/invasion of host cells (9–11). Ail and LPS mutants exhibit altered sensitivity to human serum, antibiotics and cell wall stress (12, 13) and we have identified LPS-recognition motifs on the surface of Ail important for establishing mutually reinforcing Ail-LPS interactions that promote microbial survival in human serum, antibiotic resistance and cell envelope integrity (14). These features highlight the role of the outer membrane environment as a critical regulator of Ail activity and underscore the importance of characterizing the native Ail-membrane assembly as a whole, rather than its individual components.

Previous structural studies have relied on purified Ail refolded in detergent micelles, detergent-free lipid nanodiscs or liposomes for X-ray diffraction and NMR (15–19). The structure of Ail – an eight-stranded β-barrel with four extracellular loops and three intracellular turns – is conserved in all cases, but these chemically defined samples preclude NMR studies aimed at probing the molecular interactions of Ail with either bacterial membrane components like LPS, or human ligands, in the context of the native environment. Importantly, this situation negates a major advantage of NMR compared to other structure determination technologies: the high sensitivity of NMR signals to their local environment, which makes them extremely useful for characterizing even weak inter-molecular interactions by direct spectroscopic detection.

We have shown that nanodiscs and liposomes can incorporate Ail with various types of LPS, including native *Y. pestis* LPS (14, 19), but even though these detergent-free lipid platforms represent important advances in membrane complexity they remain distant substitutes for the native outer membrane, which is highly anisotropic, heterogeneous, and connected to the cytoskeletal peptidoglycan layer. The emerging field of *insitu* NMR (20–26) presents new opportunities for examining the structural properties and interactions of Ail in a native bacterial outer membrane.

Here we describe an *E. coli* model system where Ail is natively folded in the bacterial outer membrane for parallel solid-state NMR structural studies and microbial functional assays, *in-situ*. Using this model, we demonstrate that Ail expression produces *Y. pestis* virulence phenotypes in *E. coli*, and that isolated cell envelopes expressing ^15^N and ^13^C labeled Ail yield atomic-resolution NMR spectra that allow us to probe the structure of Ail in a native bacterial membrane and its interactions with human serum components. This model sheds light on the interactions of Ail with components of human serum and provides a platform for advancing structure-activity NMR studies of Ail in the native environment, including its various physiological modulations that can be elicited by factors such as temperature, antibiotics, or exposure to serum.

## MATERIALS AND METHODS

### Expression of Ail in the *E. coli* outer membrane

The gene encoding mature *Y. pestis* Ail was cloned in the *NcoI* and *XhoI* restriction sites of the *E. coli* plasmid pET-22b(+), downstream of the signal sequence of pectate lyase B (27). This plasmid was transformed into *E. coli* cells by heat shock and positive clones were selected by plating on LB-agar with ampicillin (100 μg/mL) and chloramphenicol (35 μg/mL), in the case of Lemo21(DE3) cells. Three *E. coli* cell strains derived from BL21(DE3) (28) were tested: C41(DE3) (29), BL21(DE3)ΔACF (30) and Lemo21(DE3) (31). Transformed cells were cultured overnight, at 30°C, with vigorous shaking, in 50 mL of M9 minimal media supplemented with Basal Medium Eagle vitamin solution (1% by vol.), ampicillin (100 μg/ml), and chloramphenicol (50 μg/mL). This overnight culture was used to inoculate 500 mL of fresh M9 media (supplemented as above), and grown from OD_600_ ~0.05 to OD_600_ ~0.4, at 30°C, at which point the cells were harvested by low-speed centrifugation (5,000 g, 4°C, 20 min) and then resuspended in 500 mL of fresh supplemented M9 media. Protein expression was induced by adding 0.4 mM isopropyl β-D-1-thiogalactopyranoside (IPTG), after which the shaking speed was reduced, and the temperature lowered to 25°C. The cells were cultured for 20 min before adding rifampicin (100 μg/mL), and then for an additional 20 hours, in the dark. The cells were harvested by low-speed centrifugation, and resuspended in HEPES buffer (10 mM, pH 7.4).

To obtain uniformly ^15^N,^13^C labeled Ail, the cells were grown in unlabeled M9 media and only transferred to ^15^N,^13^C labeled M9, prepared with (^15^NH_4_)_2_SO_4_ (1 g/L) and ^13^C_6_-glucose (5 g/L), before induction with IPTG. To obtain ^2^H labeling of endogenous *E. coli* biomolecules, cells were grown in 100 mL of ^2^H_2_O and then switched to H_2_O before induction with IPTG.

### Isolation of bacterial membrane fractions

Ail-expressing cells were lysed by three passes through a French Press. After removing cellular debris by centrifugation (20,000 g, 4°C, 1 h), the total cell envelope fraction, including inner and outer membranes, was harvested by ultracentrifugation (100,000 g, 4°C, 1 h), washed three times with sodium phosphate buffer (20 mM, pH 6.5), and then collected by ultracentrifugation. To further isolate the outer membrane fraction, the total cell envelope was suspended in 7 mL of buffer I (10 mM HEPES, pH 7.4, 3.4 mM N-lauroylsarcosine) and then separated by ultracentrifugation. The supernatant containing detergent-solubilized inner membrane was removed, while the pellet was washed three times with sodium phosphate buffer (20 mM, pH 6.5) and then collected by ultracentrifugation. All three fractions – total cell envelope, inner membrane and outer membrane – were analyzed by polyacrylamide gel electrophoresis (PAGE) in sodium dodecyl sulfate (SDS), and immune-blotting with the α-Ail-EL2 antibody (17).

### NMR sample preparation

Samples for solid-state NMR studies were packed into 4 mm, 3.2 mm or 1.3 mm magic angle spinning (MAS) rotors. For samples incubated with human serum (NHS or HIS), isolated cell envelopes expressing ^13^C,^15^N Ail were suspended in 7 mL of serum and gently mixed for 4 h, at room temperature, before harvesting and washing with sodium phosphate buffer by ultracentrifugation and packing into the MAS rotor. The preparation of Ail liposome samples has been described (18).

### Solid-state NMR spectroscopy

Solid-state NMR experiments with ^13^C detection were performed on a 750 MHz Bruker Avance III HD spectrometer with a Bruker 3.2 mm ^1^H,^13^C,^15^N E-free MAS probe, or a 500 MHz Bruker Avance spectrometer with a Bruker 4 mm ^1^H,^13^C,^15^N E-free MAS probe, at an effective sample temperature of 7±5°C, with a spin rate of 11,111 Hz. Typical π/2 pulse lengths for ^1^H, ^13^C and ^15^N were 2.5 μs, 2.5 μs, and 5 μs, respectively. ^1^H decoupling was implemented with the SPINAL64 sequence, with a radio frequency (RF) field strength of 90 kHz during acquisition. Two-dimensional ^15^N/^13^C NCA spectra were acquired using the SPECIFIC-CP pulse program, with contact times of 2 ms and a 70-100% ramp, for ^1^H/^15^N cross polarization (CP), and 4.9 ms for ^15^N/^13^C CP. Two-dimensional ^13^C-^13^C PDSD (proton driven spin diffusion) spectra were acquired with a ^1^H-^13^C contact time of 1 ms, a ^13^C-^13^C mixing time of 50 ms, and 20 ms of SWf-TPPM ^1^H decoupling.

Experiments with ^1^H detection were performed on a 900 MHz Bruker Avance III HD spectrometer equipped with a Bruker 1.3 mm MAS probe, operating at an effective sample temperature of 30 ± 5°C, with a spin rate of 57,000±15 Hz. Two-dimensional ^1^H/^15^N CP-HSQC spectra were acquired as described previously (18), using the MISSISSIPPI sequence (32) for water suppression.

### Bacterial cell assays

*E. coli* Lemo21(DE3) cells transformed with Ail-expressing plasmid pET-22b(+) or with empty plasmid were cultured as described above, then harvested by low-speed centrifugation, washed twice in ice-cold phosphate buffer saline (PBS), and resuspended in ice-cold PBS to OD_600_=1.0.

To assay binding to human Vn, washed bacteria (250 μl) were added to an equal volume of either NHS (Sigma-Aldrich H4522), HIS (Sigma-Aldrich H3667), purified full-length Vn (1 μM in PBS; Sigma-Aldrich SRP3186), or purified Vn-HX (2 μM in PBS) prepared as described (33). The binding reactions were incubated for 30 min at 37°C, then placed on ice. Bacterial cells and bound proteins were then collected by centrifugation (6,000 g, 5 min, 4°C), washed three times with 1 mL of ice-cold PBS containing 0.1% Tween-20, and lysed by boiling in 100 μL of SDS-PAGE sample buffer (50 mM Tris-HCl, 2% SDS, 5% glycerol, 1% β-mercaptoethanol, pH 6.8). Bacterial cell lysates and co-sedimented proteins were subjected to SDS-PAGE and immunoblot analysis with rabbit polyclonal anti-Vn (R12-2413, Assay Biotech) and anti-Ail-EL2 (34) antibodies.

To assay serum resistance, *E. coli* cells collected after incubation with NHS or HIS were diluted 100-fold with PBS, and 25 μL of this dilution mixture were plated on LB-agar supplemented with ampicillin (100 μg/mL) and chloramphenicol (35 μg/mL). After incubating overnight at 37°C overnight, the percentage of cell survival was estimated by counting the number of bacterial colonies present on each plate, using ImageJ (35). The percent survival represents the number of colonies that survive in NHS divided by the number that survive in HIS.

To assay pellicle formation, *E. coli* cells were harvested, then resuspended in 2 mL of M9 minimal media to OD_600_ = 0.5 in 15 x 100 mm glass tubes, and incubated for 16 h at 37°C. After removing the cells by centrifugation, the interior glass walls were treated with methanol and air-dried overnight, then washed three times with PBS and again air-dried overnight. The tubes were treated with 2.5 mL of crystal violet solution (0.1 %) for 10 min, then washed with PBS and air-dried. Pellicle formation was detected qualitatively as a violet-stained rim at the air-liquid interface.

## RESULTS AND DISCUSSION

### Production of folded Ail in the outer membrane of *E. coli*

High-resolution, multi-dimensional NMR studies require isotopically ^15^N and ^13^C labeled biomolecules. For studies of proteins in native cell membranes, targeted isotopic labeling is critical for suppressing background NMR signals from other cellular components, and solidstate NMR methods are needed to overcome the correlation time limitations posed by samples that are effectively immobilized on a time scale ≥ μsec. Moreover, the inherently low sensitivity of NMR necessitates samples that are enriched in target protein. This requirement is compatible with the properties of Ail, which naturally comprises more than 30% of the *Y. pestis* outer membrane proteome at the mammalian infection temperature of 37°C (6, 9, 36–38).

*E. coli* is widely used as a model for assaying the functions of *Y. pestis* proteins, including Ail. *E. coli* strains derived from BL21(DE3) have a rough-type LPS that lacks an extended O-antigen, similar to that of *Y. pestis,* and we have shown that *E. coli* rough-type LPS and *Y. pestis* LPS both perturb the NMR spectra of Ail in a similar manner (14). The use of *E. coli* also facilitates isotope labeling for NMR.

To drive the production of folded, ^15^N,^13^C-Ail in the *E. coli* outer membrane, we cloned the sequence of the pectate lyase B (PelB) leader peptide (27) before the N-terminus of mature Ail, and relied on the *E. coli* native β-barrel assembly machinery (BAM) to insert Ail across the bacterial outer membrane. The Ail-enriched outer membrane or the entire cell envelope – including inner and outer membranes -were isolated and taken directly for NMR (Fig. 1A). To reduce isotopic labeling of endogenous *E. coli* components, the cells were grown in unlabeled minimal media and transferred to isotopically labeled media only before induction with IPTG. We tested Ail expression in three derivative strains of *E. coli* BL21(DE3) (28), where recombinant protein synthesis is controlled by a chromosomally-encoded, IPTG-inducible polymerase from bacteriophage T7 and its plasmid-encoded T7 promoter.

**Figure 1.**
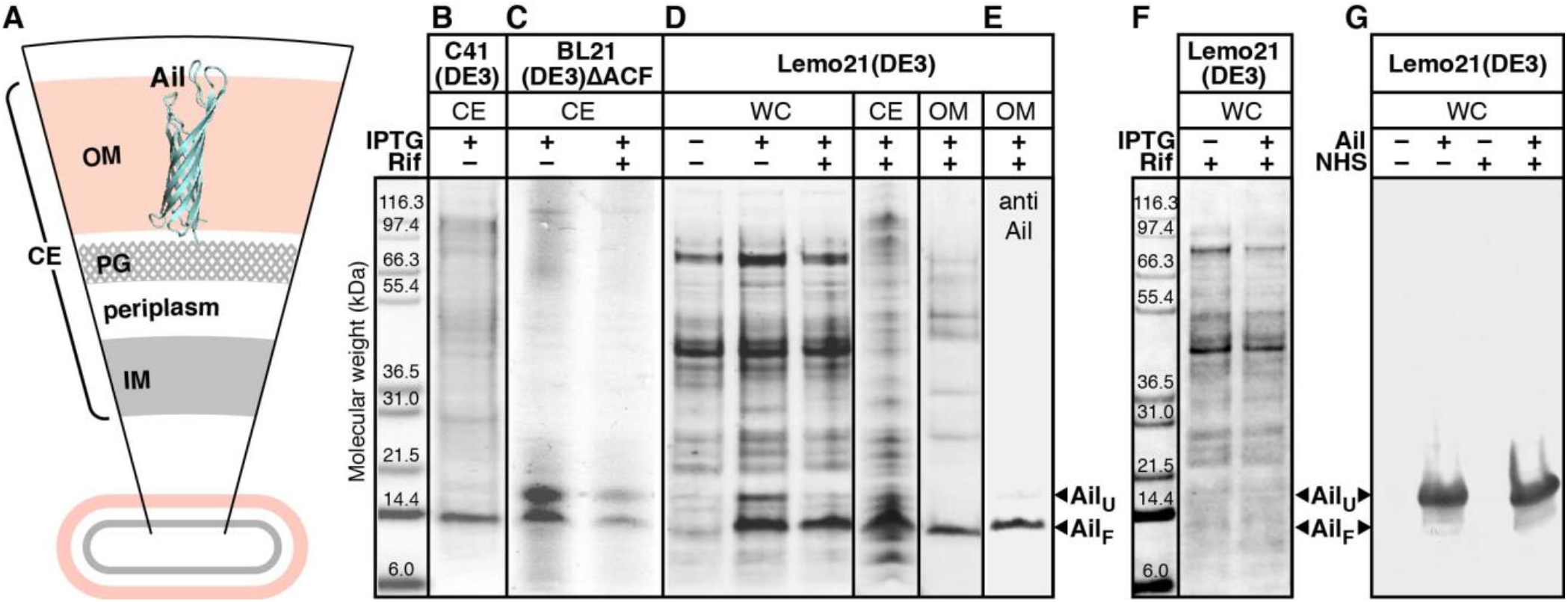
Production of folded Ail in *E. coli* and in situ NMR of Ail in *E. coli* cell envelopes. **(A)** Depiction of the bacterial cell envelope (CE) fraction isolated for NMR studies, including the outer membrane (OM) with embedded Ail (cyan), inner membrane (IM), peptidoglycan layer (PG), and periplasm. **(B-F)** SDS-PAGE analysis of cell envelope (CE), whole cell (WC), or outer membrane (OM) fractions isolated from *E. coli* C41(DE3), BL21(DE3)ΔACF, or Lemo21(DE3) cells transformed with Ail plasmid (B-E) or empty plasmid (F). Cells expressing Ail (+) or empty plasmid (-) were incubated with NHS for 30 min and then analysed by SDS-PAGE and immunoblotting after heat denaturation (G). Cells were grown with or without rifampicin and induced with IPTG. Ail was visualized with Coomassie stain (B-D, F) or immunoblotting with Ail-specific antibody (E, G).

Our initial attempts utilized C41(DE3) cells (29), which promote folded protein insertion in the *E. coli* membrane due to a mutation that weakens the production of T7 polymerase (31). These cells yield high expression of properly folded Ail (Fig. 1B), but also produce appreciable amounts of endogenous *E. coli* proteins that are isotopically labeled. Background NMR signal can be reduced by expressing Ail in BL21(DE3)ΔACF, a mutant cell strain that lacks the major *E. coli* outer membrane proteins OmpA, OmpC and OmpF (30). Compared to C41 strains, this mutant is compatible with the use of rifampicin, a potent inhibitor of *E. coli* RNA polymerase (39) that suppresses endogenous protein production was first used to enable targeted isotopic labeling of heterologous proteins for solution (40) NMR studies *in situ.* These cells produce very high levels of Ail (Fig. 1C), but their usefulness is limited by the production of large amounts of misfolded protein, detectable as a band migrating just above 17 kDa in sodium dodecyl sulfate polyacrylamide gel electrophoresis (SDS-PAGE) that is distinguishable from folded Ail at 14 kDa (17). The addition of rifampicin to BL21(DE3)ΔACF cells reduces protein expression levels overall, but does not enhance the proportion of folded to unfolded Ail. Although endogenous background signal can be reduced by isolating the outer membrane from the cell envelope, this process exposes the sample to the compromising effects of detergent on biomolecular structure and activity (41, 42). The two-dimensional (2D) ^15^N/^13^C NCA solid-state NMR spectra from outer membrane preparations of either C41(DE3) or BL21(DE3)ΔACF cells reflect the overall signature of Ail (17–19) but suffer from sub-optimal resolution and sensitivity (Fig. S1).

To avoid these complications we tested Lemo21(DE3) cells (31), where T7 RNA polymerase can be controlled by its natural inhibitor T7 lysozyme, and rifampicin can be used to effectively halt transcription of the *E. coli* genome. By initially growing the cells in unlabeled minimal media, then transferring them to isotopically labeled media supplemented with both rifampicin and IPTG, expression of ^15^N/^13^C-Ail was induced and allowed to continue under control of the bacteriophage T7 promoter, while endogenous protein production was blocked. As an additional important advantage, we found that rifampicin moderates the level of Ail overexpression and reduces protein misfolding to negligible levels, resulting in cell envelope and outer membrane fractions that are highly enriched in folded Ail (Fig. 1D, E).

### Solid-state NMR of Ail in bacterial cell envelopes

Cell envelope preparations from Lemo21(DE3) cells yield high quality 2D ^15^N/^13^CA solid-state NMR spectra, with respect to both signal intensity and resolution (Fig. 2A, black). The line widths of well-resolved signals, such as those from Ala and Gly (Fig. S2), are in the range of 0.6-0.8 ppm for ^13^C and 1.3-1.8 ppm for ^15^N. They compare favorably with those measured from purified Ail reconstituted in liposomes (18), demonstrating that the combined use of this cell strain with rifampicin is highly beneficial for producing folded, isotopically labeled Ail. Based on comparison of the 1D cross-sections of the ^15^N/^13^C peaks (Fig. S2, S6) with those from Ail-liposomes (18), we estimate that the cell envelope sample in the 3.2 mm MAS rotor contains approximately 0.7 mg of Ail or 25% of the Ail-liposomes.

**Figure 2.**
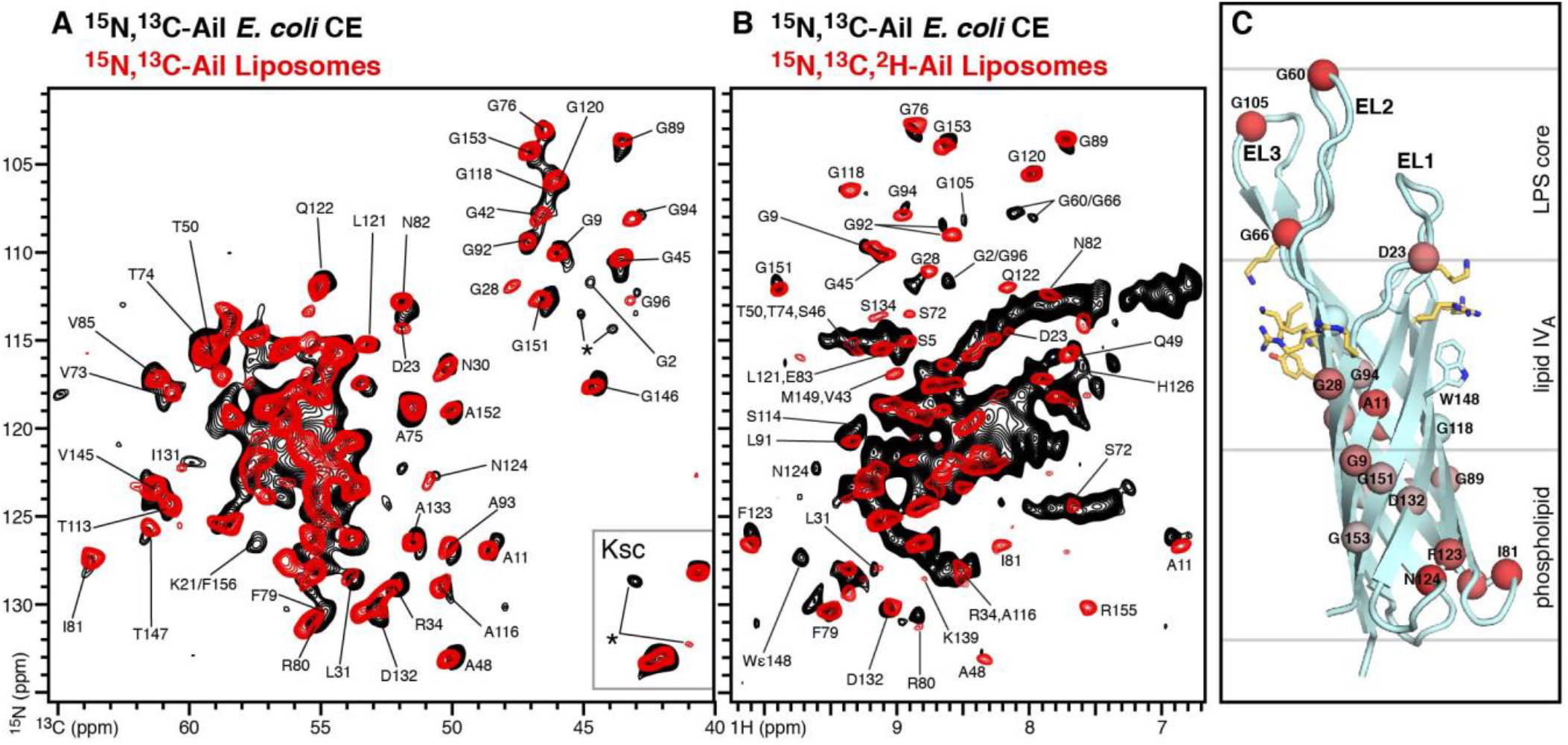
Solid-state ^15^N/^13^C/^1^H NMR spectra of Ail in *E. coli* cell envelopes. **(A)** 2D ^15^N/^13^C NCA spectra of ^15^N,^13^C-Ail in *E. coli* cell envelopes (black) or reconstituted liposomes (red). Spectra were recorded at 750 MHz, 7°C, with a MAS rate of 11 kHz and 640 transients for the cell envelope samples or 128 transients for the liposome samples. Resolvable assigned peaks are marked. Asterisks denote new unassigned peaks. **(B)** ^1^H-detected ^1^H/^15^N CP-HSQC spectra of ^15^N-Ail in *E. coli* cell envelopes (black) or in reconstituted liposomes (red). Spectra were recorded at 900 MHz, 30°C, with a MAS rate of 57 kHz and 1,600 transients for the cell envelope sample or 160 transients for the liposome samples. **(C)** Structural model of Ail embedded in the *Y. pestis* outer membrane taken from previous MD simulations (14). Spheres denote resolved and assigned amide N atoms that undergo ^1^H/^15^N chemical shift perturbations from 0 ppm (cyan) to 0.15 ppm (red), between the cell envelope and liposome environments. Sidechains forming two clusters of LPS-recognition motifs are shown as yellow sticks. The boundaries of the outer membrane phospholipid and LPS layers are marked (gray lines). Residue numbers, from E1 to F156, corresponds to the mature sequence of Ail.

Notably, the 1D ^13^C spectra (Fig. S3) acquired for *E. coli* cell envelopes prepared from Ail(-) bacteria, transformed with empty plasmid but otherwise grown in an identical manner as Ail(+) bacteria, show no evidence of protein signals when compared to the spectra of Ail(+) preparations. These cells do not express Ail (Fig. 1F) and, together with the NMR data, they demonstrate that the expression system is tightly regulated and leads to highly selective isotope labeling of the target Ail, with minimal background.

Many resonances in the 2D ^15^N/^13^CA spectrum can be assigned by direct comparison with the solid-state and solution NMR spectra of purified Ail reconstituted in liposomes (Fig. 2A, red) or nanodiscs (18, 19), indicating that the same overall structure – an eight-stranded β-barrel with four extracellular loops (EL1-EL4) and three intracellular turns (T1-T3) (15, 16, 19) – is preserved in the bacterial outer membrane. Within the 2D ^15^N/^13^CA spectrum, there are some new signals and 47 cross-peaks are sufficiently resolved to facilitate the detection of site-specific chemical shift perturbations (relative to liposomes) for residues dispersed across the Ail sequence and its β-barrel topology (Fig. S4A).

The ^1^H, ^15^N and ^13^C chemical shifts from backbone sites are sensitive to chemical environment, ligand binding and protein conformational changes (43). To examine the effects of the native membrane environment on Ail, we acquired a ^1^H-detected ^1^H/^15^N cross polarization (CP) solid-state NMR spectrum of Ail in bacterial cell envelopes (Fig. 2B, black). Typically, ^2^H labeling of side chain protons is needed to achieve line narrowing and high sensitivity in ^1^H-detected solid-state NMR experiments with MAS rates ~60 kHz (44, 45). For purified and reconstituted membrane proteins, this can be achieved by labeling with ^2^H and back-exchanging the amide hydrogens to ^1^H during reconstitution with lipids. This approach, however, is not readily applicable to cellular membrane proteins where membrane-embedded segments remain water-inaccessible throughout the sample preparation protocol. In this study, we prepared the sample by initially growing cells in ^2^H_2_O, and then transferring the culture to ^1^H_2_O, with ^15^N salts and rifampicin, for Ail induction. The goal was to silence the background by ^2^H labeling (~70%) of endogenous *E. coli* components, whilst achieving targeted ^15^N labeling of Ail. The resulting 2D CP-HSQC spectrum is strikingly good, despite the lack of Ail deuteration.

The cell envelope spectrum, obtained at 900 MHz ^1^H frequency and 30°C, compares favorably with the CP-HSQC solid-state NMR spectrum of fractionally deuterated (~70%) Ail in liposomes (Fig. 2B, red), which was obtained at the same field and temperature (18), and also with the TROSY-HSQC solution NMR spectrum of uniformly deuterated (~99%) Ail in nanodiscs, obtained at 800 MHz and 45°C (Fig. S5) (18, 19). Individual resonances have line widths in the range of 0.15 ppm for ^1^H and 1.5 ppm for ^15^N, in line with recent reports of ^1^H-detected spectra of membrane proteins in native cell membranes (46). A total of 22 ^1^H/^15^N peaks can be resolved and assigned by direct comparison with the previously assigned solution NMR spectra of nanodiscs and solid-state NMR spectra of liposomes (18, 19). Improvements in ^1^H/^15^N resolution should be available through the implementation of solid-state NMR experiments with MAS rates greater than 60 kHz as described (44, 45), and ^2^H labeling schemes such as those that enable selective visualization of the water-accessible loops, obtained by growing cells in ^2^H-labeled media followed by incubation in H_2_O to back-exchange water-accessible sites.

Previously, we showed that Ail acquires conformational order in the presence of LPS, leading to substantial enhancement of solid-state NMR CP signal intensity (19). Moreover, all-atom MD simulations (14) indicate that the extracellular loops of Ail exhibit dramatically reduced root mean square fluctuations in a native bacterial membrane. In line with these earlier observations, the NCA and CP-HSQC spectra of Ail in cell envelopes contain some new signals, which were not previously observed in the spectra from reconstituted liposomes. These include HN peaks assigned in the solution NMR spectra of nanodiscs (G2, G60, G66, G105, S114, and the side chain of W148), as well as new unassigned peaks. These results suggest that Ail has more restricted conformational dynamics in *E. coli* outer membranes leading to enhanced CP transfer efficiency and signal intensity.

Spectral comparisons reveal chemical shift differences between the chemically defined liposome membrane environment and the bacterial cell envelope (Fig. 2C; Table S1). Independent spectroscopic assignments will be needed to fully and precisely map the effects of the bacterial membrane environment on the protein, but some prominent differences are detectable at this stage. These map to sites in the extracellular loops (D23, G60, G66, G105) and near the extracellular membrane-water interface (A11, G28, A48, G92, G94), just below the two clusters of positively charged residues that we have identified as LPS-recognition motifs on opposite poles of the Ail β-barrel (14): cluster I (R14, K16, K144 and K139), which is tightly localized to the base of the two short extracellular loops EL1 and EL4, and cluster II, which occupies a broader region on the barrel surface extending from the base (R27, R51, H95) to the outer extremities (K69, K97, K99) of the two long loops EL2 and EL3.

Notable spectral differences are also observed for membrane-embedded sites of the Ail β-barrel (G9, G89, D132,G151, G153) and sites in the intracellular turns (R80, I81, F123, N124), suggesting that the entire protein senses the physical-chemical properties of its membrane environment. The results are in line with our recent report (14) that specific interactions of outer membrane LPS with cluster I and cluster II Ail sites enhance the conformational order of Ail, dampen LPS dynamics and cause an overall thickening of the outer membrane around its β-barrel, while mutations in the Ail binding sites for LPS compromise cell envelope integrity, reduce *Y. pestis* survival in serum, and enhance antibiotic susceptibility.

To further examine the side chain conformations of Ail in the bacterial cell envelope, we acquired a 2D ^13^C/^13^C correlation PDSD spectrum (Fig. 3 black). The spectrum, obtained in 25 hours, has excellent resolution. The overall pattern of cross peaks is conserved between bacterial cell envelope and liposome preparations (Fig. 3, black), indicating that the side chain conformations are broadly similar. Resolved signals from Ala, Ile, Pro, Ser, Thr and Val spin systems can be easily identified based on their chemical shifts, which reflect the β-barrel structure of Ail, and 54 intra-residue cross-peaks can be assigned (Fig. S6) by direct transfer from the liposome NMR data which was assigned previously (18). By contrast, the spectrum of Ail(-) cell envelopes contains no protein signals (Fig. S6), confirming the isotopic labeling selectivity of the expression system.

**Figure 3.**
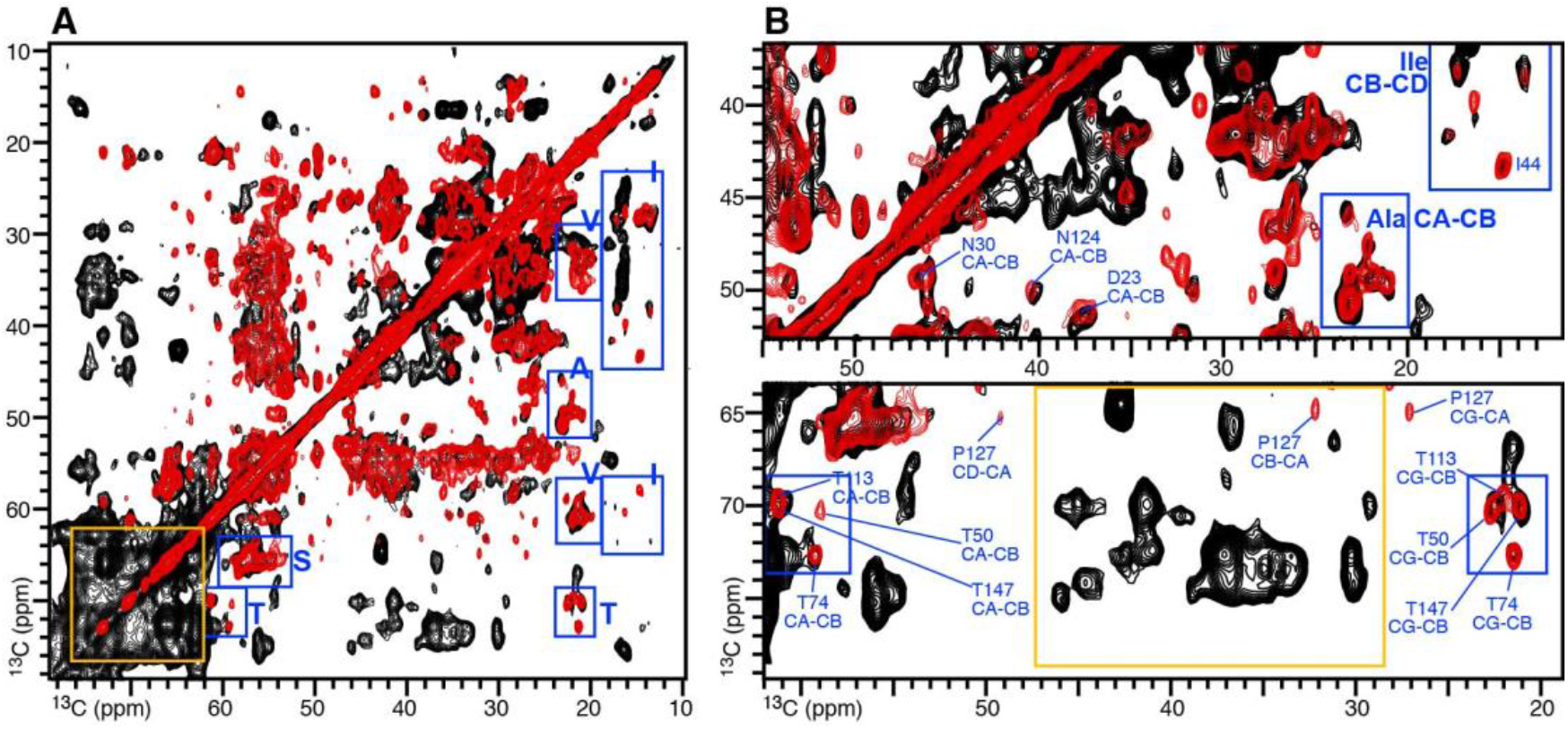
Solid-state ^13^C/^13^C PDSD NMR spectra of ^15^N,^13^C-Ail in *E. coli* cell envelopes. Resolvable signals from Ala Ile, Ser, Thr, and Val residues of Ail (blue) and non-Ail bacterial cell envelope components (gold) are marked. **(A)** Aliphatic region of the 2D spectra from *E. coli* cell envelopes (black) or reconstituted liposomes (red). Spectra were recorded at 750 MHz, 7°C, with a MAS rate of 11 kHz, 204 t1 increments and 304 transients for the cell envelope sample, or 512 t1 increments and 64 transients for the liposome sample. **(B)** Expanded spectral regions.

The ^13^C/^13^C correlation PDSD spectrum also contains signals from cell envelope components other than Ail, including lipids, LPS and peptidoglycan, which make up a large proportion of the bacterial cell mass and incorporate ^13^C even with the use of rifampicin, due to relatively fast doubling time of *E. coli* cells and the lag time between IPTG induction and rifampicin addition. While the flexible regions of these molecules are detectable only by through-bond polarization transfer, their more rigid groups are visible in the NMR spectra obtained with CP (23, 47) and are present in the PDSD spectra of both Ail(+) and Ail(-) cell envelopes (Fig. S6).

Treatment with cerulenin, an inhibitor of fatty acid biosynthesis has been used to inhibit ^13^C labeling of *E. coli* lipids and simplify ^13^C/^13^C correlation spectra of proteins (22). Moreover, spectroscopic approaches have been developed to silence lipid signals based on ^15^N-filtering (48), and NMR data acquisition above the gel-to-liquid phase transition of the lipids has also been shown to suppress lipid signal intensity (49). These cell envelope signals, however, also present an opportunity to probe specific interactions of Ail with the cell envelope.

Notably, a set of ^13^C/^13^C cross peaks observed in the region between 30-45 ppm and 60-75 ppm of the Ail(+) cell envelope spectrum are absent from the Ail(-) spectrum. Resonance assignments will be needed to determine the network of inter-atomic interactions in the Ail outer membrane, but there are two plausible explanations for this apparently Ail-dependent observation. Firstly, these resonances could reflect contacts between LPS sugar moieties (60-75 ppm) and His, Lys, Arg and Tyr sidechains of Ail (30-45 ppm) in line with previous observations based on chemical shift perturbation analysis (14). Secondly, they could reflect inter- or intra-molecular contacts of LPS and lipids, resulting from the general ordering effect of Ail on the outer membrane of *E. coli*. These effects need not be exclusive but could synergize such that Ail-LPS interactions rigidify bound LPS molecules thereby enhancing LPS intra-molecular CP transfer.

### Emergence of *Y. pestis* phenotypes in *E. coli* through Ail expression

One important advantage of *in-situ* NMR spectroscopy is the ability to probe the structural underpinnings of biology by assaying protein activity in the same samples that are analyzed by high resolution spectroscopy. Having confirmed the structural integrity of Ail expressed in *E. coli* cells, we asked whether Ail also presents its characteristic virulence phenotypes observed in *Y. pestis*.

The role of Ail in providing serum resistance to *Y. pestis* is well known, and expression of Ail in *E. coli* DH5α cells is sufficient to render them immune to serum mediated killing (34). Similarly, we found that Ail(+) Lemo21(DE3) cells, expressing plasmid-encoded Ail, exhibit 87 ± 7% survival after incubation with normal human serum (NHS) relative to heat-inactivated serum (HIS). By contrast, Ail(-) cells, transformed with empty plasmid, formed no colonies after incubation with NHS (Fig. 4A).

**Figure 4.**
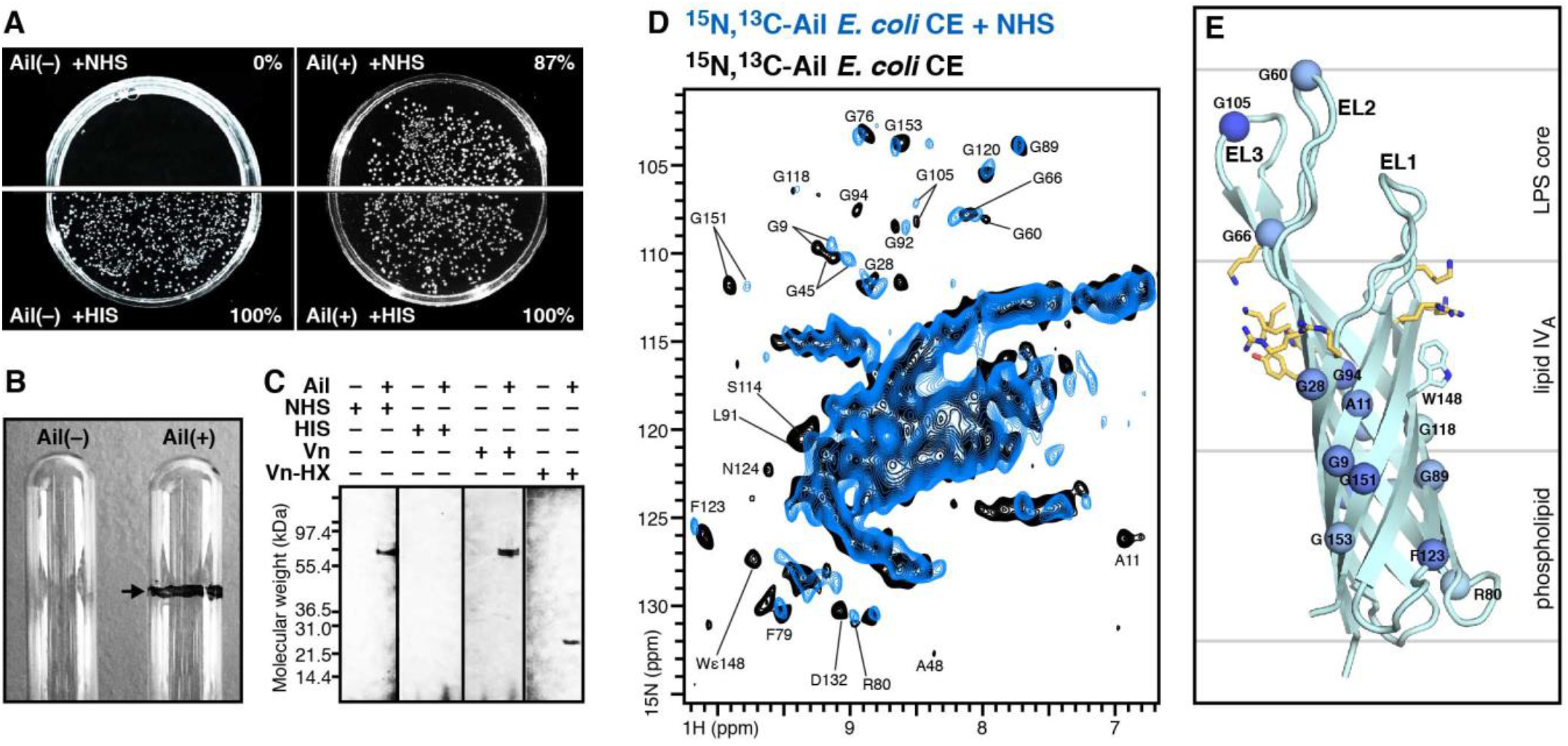
*Y. pestis* phenotypes induced in *E. coli* by Ail expression and interaction of Ail-expressing *E. coli* cells with human serum. Assays were performed with *E. coli* Lemo21(DE3) cells transformed with Ail-bearing plasmid, Ail(+), or with empty plasmid, Ail(-). Each data set is representative of at least triplicate experiments. **(A)** Survival of *E. coli* cells in normal human serum (NHS) relative to heat-inactivated serum (HIS) was assayed by incubating cells with either serum, then plating on agar and counting the surviving bacterial colonies. Survival is reported as percent serum resistance relative to the number of surviving colonies in HIS. **(B)** Pellicle formation observed in Ail(+) but not Ail(-) cells. Cells were suspended in 2 mL of M9 minimal media (OD_600_ = 0.5, in 15 x 100 mm glass tubes) and incubated for 16 h at 37°C. After removing the cells by centrifugation, the interior glass walls were treated with methanol and air-dried overnight, then washed three times with buffer and air-dried overnight. Finally, the tubes were treated with 2.5 mL of crystal violet solution (0.1% for 10 min) then washed with buffer and air-dried. Pellicle formation is detected as a violet-stained rim at the air-liquid interface (arrow). **(C)** Vn binding activity assayed by co-sedimentation of Ail(+) or Ail(-) bacterial cells, with NHS, HIS, purified full-length Vn, or purified Vn-HX. Immunoblots were probed with anti-Vn or anti-Ail antibodies. **(D)** ^1^H-detected ^1^H/^15^N CP-HSQC solid-state NMR spectra of Ail in *E. coli* cell envelopes acquired before (black) or after (blue) incubation with NHS. Spectra were recorded at 900 MHz, 30°C, with a MAS rate of 57 kHz, and 1,600 (black) or 2048 (blue) transients. **(E)** Structural model of Ail embedded in the *Y. pestis* outer membrane taken from a recent MD simulation (14). Spheres denote resolved and assigned amide N atoms that undergo ^1^H/^15^N chemical shift perturbations from 0 ppm (cyan) to 0.20 ppm (blue), in the presence of NHS. The boundaries of the outer membrane phospholipid and LPS layers are marked.

Ail is also known to promote bacterial cell auto-aggregation, pellicle formation and flocculent bacterial growth (50), three phenotypes that are associated with virulence in *Y. pestis* strains and many other microbial pathogens. In the case of Ail, this property is thought to be related to the distribution of charged amino acids in its extracellular loops. Notably, Ail(+) Lemo21(DE3) cells displayed marked auto-aggregation, and formed a pellicle at the air-liquid interface that can be readily visualized by staining with crystal violet (Fig. 4B). By contrast, neither Ail(-) or native non-transformed cells formed pellicles.

Finally we tested the ability of Ail(+) Lemo-21(DE3) cells to bind known ligands of Ail. Previous studies have shown that *Y. pestis* relies on Ail to associate with many human host proteins, including fibronectin (51), C4b-binding protein (52), and vitronectin (Vn) (33, 34). Incubation of Ail(+) cells with either NHS, purified fulllength Vn, or purified Vn-HX – corresponding to the hemopexin-like (HX) domain of Vn (33) – resulted in Vn co-sedimentation in all three cases (Fig. 4C). By contrast, no co-sedimentation was observed for either Ail(-) cells or Ail(+) cells incubated with HIS where Vn is presumably denatured by heat treatment (34). We conclude that Ail confers specific virulence-related *Y. pestis* phenotypes to Lemo21(DE3) cells. Notably, this implies that the samples for structural analysis are taken from a biologically active cellular system.

### Interaction of Ail with human serum components

Identifying specific Ail sites involved in serum resistance is of high biomedical importance. To reconstitute the host-pathogen interface that exists when *Y. pestis* enters the blood stream and study this microenvironment at atomic resolution, we incubated Ail(+) cell envelopes with NHS, washed extensively with buffer to eliminate non-specific binding, and then transferred the sample to the NMR rotor for spectroscopic analysis. Previous studies (34) have shown no evidence of Ail proteolysis in the presence of serum and we confirmed that this was indeed the case by immunoblot analysis of Ail after incubation with NHS (Fig. 1G; note that the sample was heat-denatured prior to SDS-PAGE and immunoblotting, thus Ail migrates at the apparent molecular weight of unfolded protein). Incubation with NHS produced several chemical shift perturbations in the ^1^H/^15^N spectrum of Ail (Fig. 4D, E; Fig. S4B).

Ail is known to bind at least two serum components C4BP (34, 52) and Vn (33, 34). While additional studies with purified Ail ligands will be needed to assign the observed perturbations to specific serum molecules, it is notable that a few ^1^H/^15^N signals are appreciably perturbed. These include signals from G66, G92, G94, residues that localize near F54, F68, S102, and F104 which have been shown to be critical for binding serum Vn and for the serum resistance phenotype (53). Interestingly, intra-membrane perturbations, near the center of the Ail transmembrane β-barrel, are also observed, indicating that the interactions of serum molecules with the extracellular loops of Ail may relay allosterically to membrane-embedded sites. Moreover, it is possible that the Ail β-barrel senses outer membrane perturbations caused by the association of serum components with the bacterial cell surface.

### Conclusions

The ability to probe the molecular structure and interactions of Ail in its native cellular environment is critical for gaining mechanistic insights into Ail-mediated pathogenesis and for developing medical countermeasures. Previously, using NMR and MD simulations with artificial membranes of defined molecular composition, we identified specific molecular interactions between Ail and LPS that are important for *Y. pestis* resistance to serum and antibiotics (14). Here, we have shown that Ail and its virulence phenotypes can be expressed in a bacterial outer membrane for parallel, *in situ* microbiology assays and solid-state NMR structural studies with atomic-resolution. These tools allow us to expand the size and complexity of the Ail microenvironment further than previously possible, and identify key residues that are sensitive to both the bacterial membrane environment and the interactions of human serum.

In NMR studies with bacterial cell envelopes, we detect specific Ail signals that firmly establish the sensitivity of the protein’s conformation to its microenvironment. Perturbations across distinct topological regions of Ail can be detected as a response to its expression in the asymmetric bacterial outer membrane, and while the overall structure of Ail is unchanged, the native membrane appears to have subtle ordering effects on the extracellular loops. The NMR data further reveal sensitivity of specific Ail sites to serum exposure, and while the identity of the serum components responsible for these effects remains to be determined, human Vn is a good candidate as it is avidly recruited by Ail to the bacterial cell surface. Finally, the NMR data indicate that Ail may sense serum-induced perturbations of the surrounding membrane environment. While the mechanism underlying these observations is unknown, the recruitment of complement components at the cell surface could be responsible for such an effect.

The present results provide a molecular window to the Ail-membrane assembly. They reaffirm the importance of the membrane environment as a key regulator of biological function and underscore the importance of characterizing the native assembly as a whole, rather than its individual components.

## Acknowledgements

This study was supported by grants from the National Institutes of Health (GM 118186, GM 122501, AI 130009 and P41 EB 002031) and a postdoctoral fellowship from Canadian Institutes of Health Research (to KS).

## Supporting Information

This article contains supporting information online.

## Author Contributions

JEK, LMF, KS, CS, YY and SP: experiment execution JEK, SJO, GVP, FMM: research design; data analysis; writing and editing

## Data Availability

All data needed are present in the paper and/or the Supplementary Materials. Additional information is available upon request.

