## Supplementary Figures and Tables for "Correlating the structure and activity of *Y. pestis* Ail in a bacterial cell envelope"

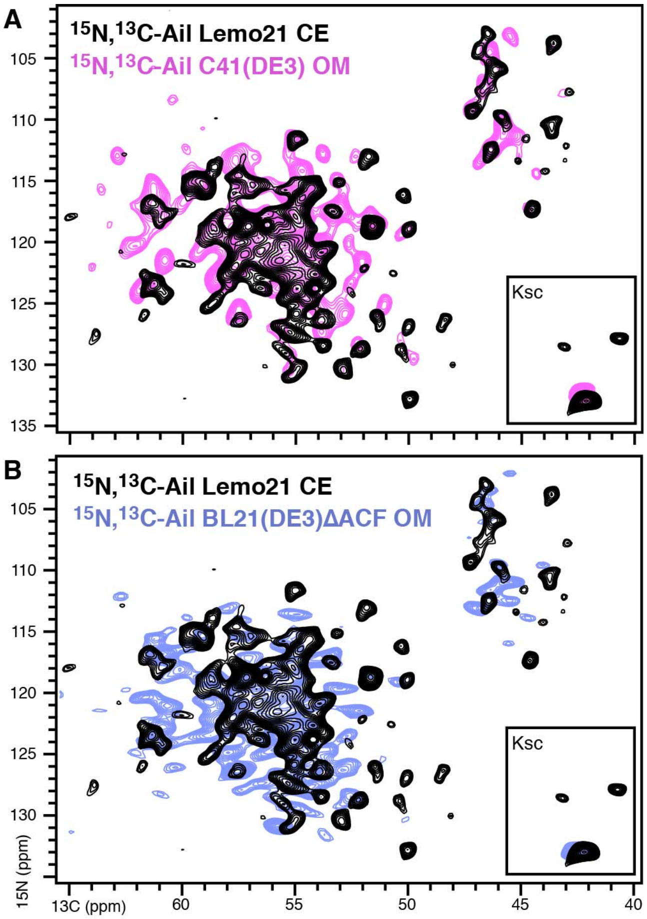


**Figure S1. 2D $^{15}$N/$^{13}$C NCA solid-state NMR spectra of Ail in *E. coli* cell envelopes (CE), *E. coli* outer membranes (OM) and liposomes. (A, B)** NMR spectra were acquired for $^{15}$N,$^{13}$C-Ail in *E. coli* CE from Lemo21(DE3) cells (black) or *E. coli* OM from C41(DE3) cells (pink), *E. coli* OM from BL21(DE3)ΔACF cells (violet). NMR spectra were recorded at 750 MHz, 7°C, with a MAS rate of 11 kHz. Inset box denotes folded-in peaks from Lys side chains (Ksc).

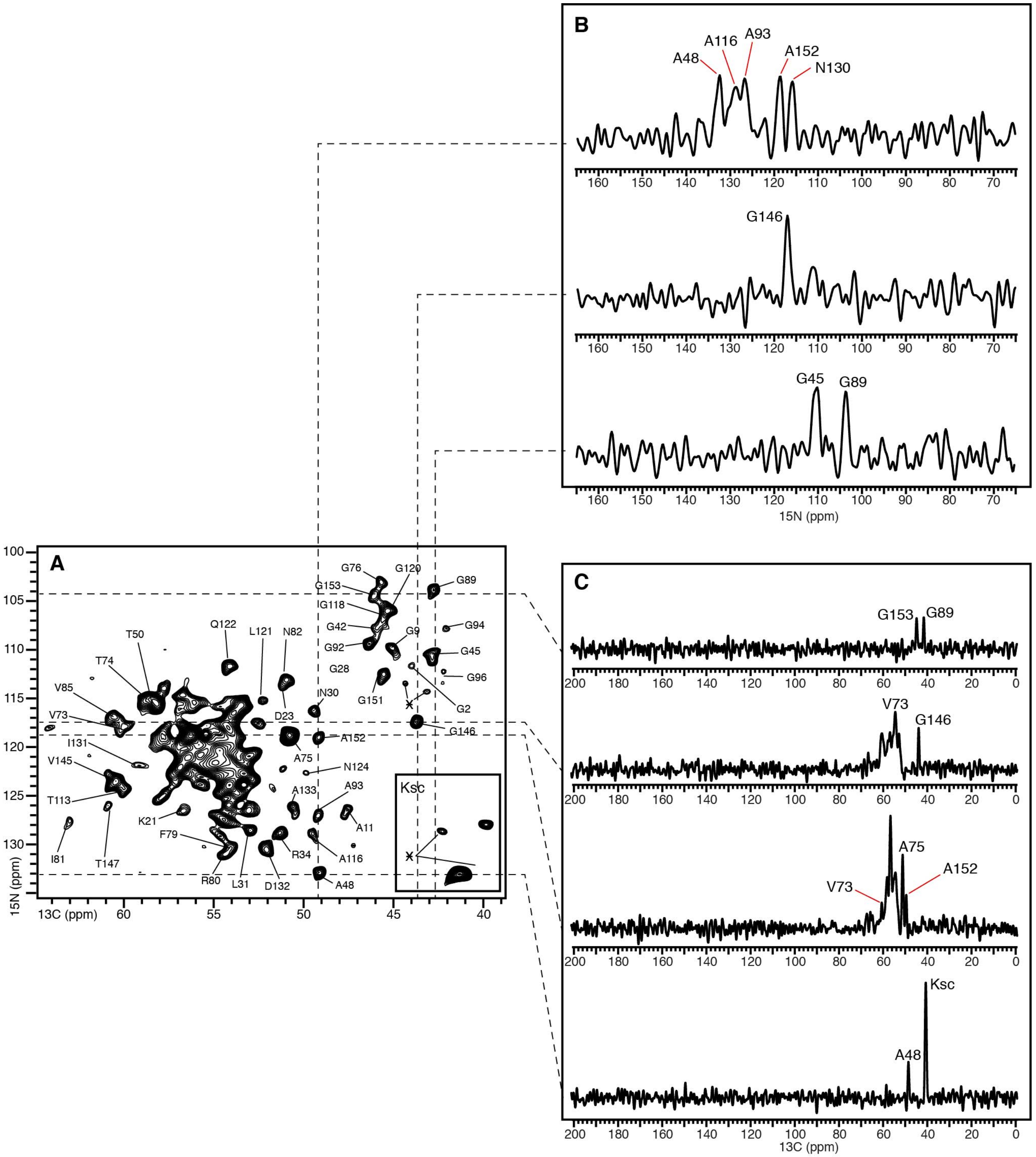


**Figure S2. Solid-state NMR $^{15}N/^{13}C$ NCA spectrum of $^{15}N,^{13}C$-Ail in *E. coli* cell envelopes.** The data were recorded at 750 MHz, 7°C, with a MAS rate of 11 kHz and 640 transients. Resolvable assigned peaks are marked. Asterisks denote new unassigned peaks. **(A)** 2D spectrum. **(B)** 1D $^{15}N$ or $^{13}C$ spectra taken from the 2D spectrum.

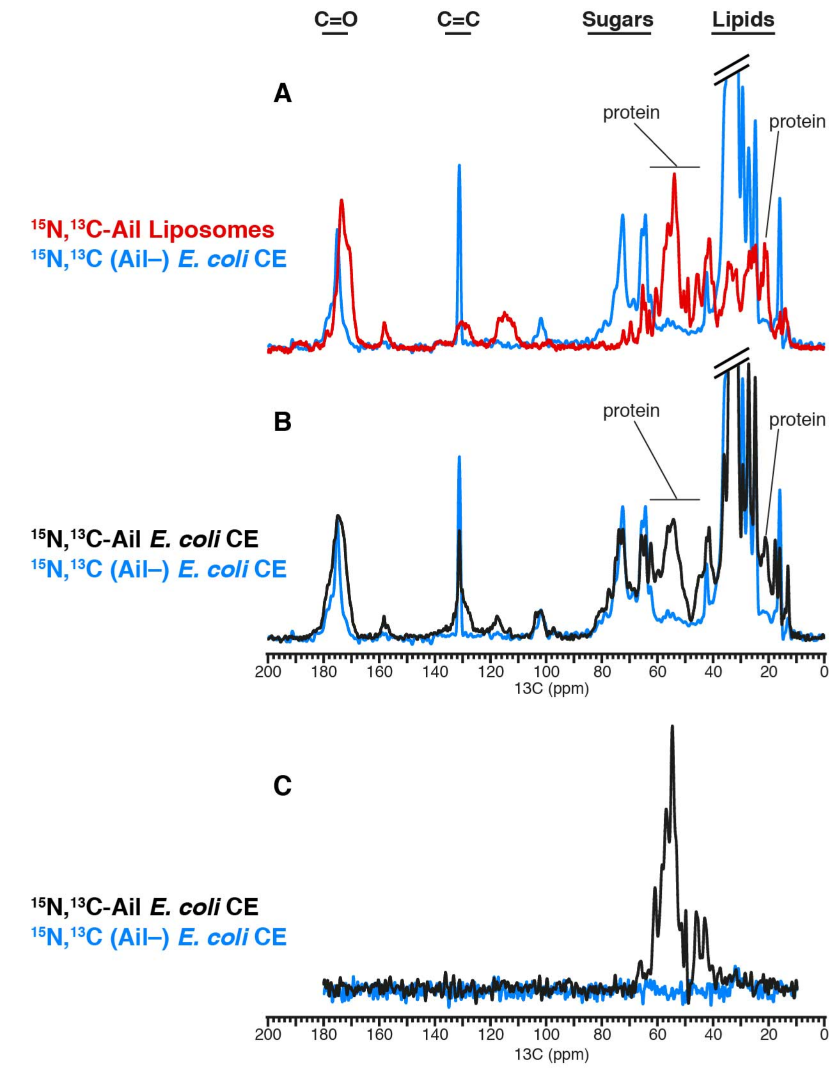


**Figure S3. 1D $^{13}C$ solid-state NMR spectra.** The spectra were obtained with cross polarization for $^{15}N$,$^{13}C$-Ail in liposomes (red), $^{15}N$,$^{13}C$-Ail in cell envelopes or *E. coli* cell envelopes (black), or $^{15}N$,$^{13}C$ Ail(–) *E. coli* cell envelopes (blue). Signals from protein and CO, CC sugar and lipid groups are marked. **(A,B)** 1D NCA spectra. **(C)** 1D CP spectra.

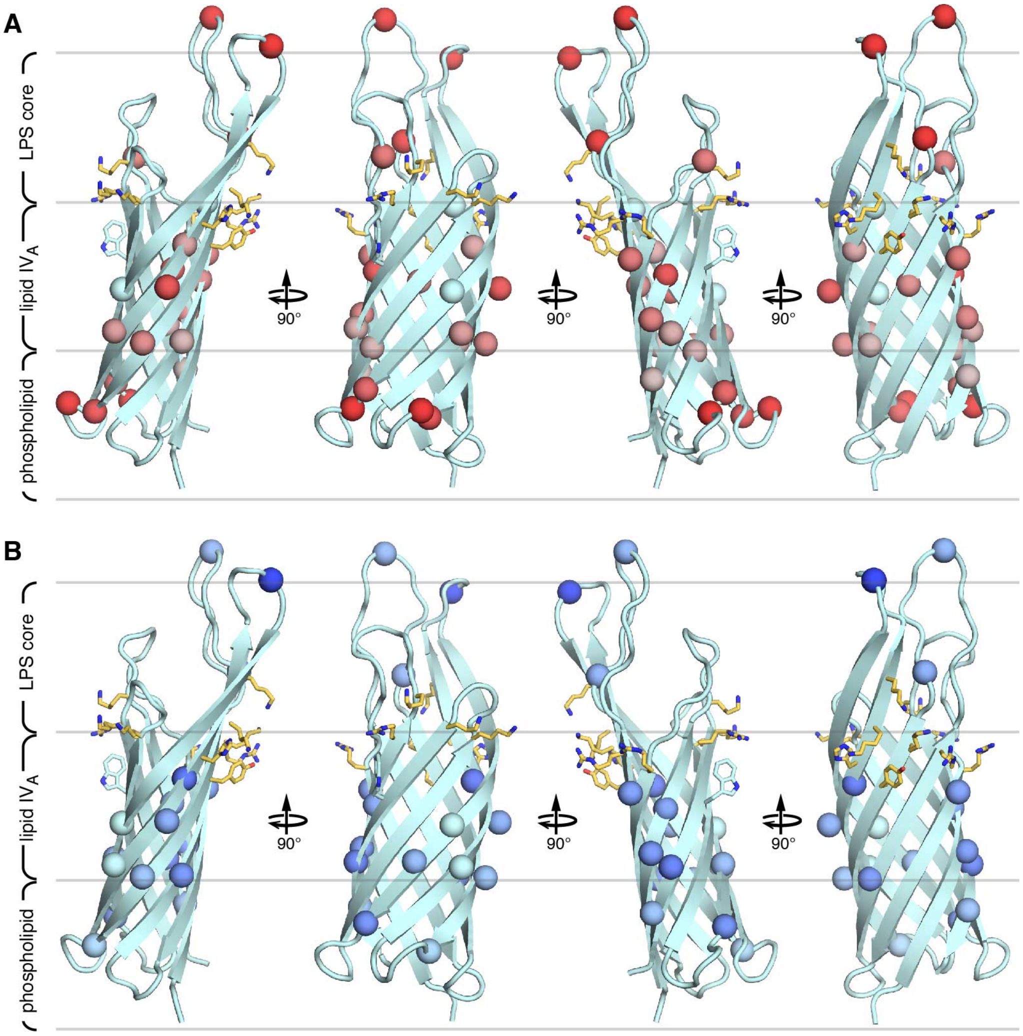


**Figure S4. Structural model of Ail embedded in the *Y. pestis* outer membrane showing perturbations associated with the cell envelope environment or interactions with NHS.** Sidechains forming two clusters of LPS-recognition motifs are shown as yellow sticks. The boundaries of the outer membrane phospholipid and LPS layers are marked (gray lines). Residue numbers, from E1 to F156, corresponds to the mature sequence of Ail. **(A)** Spheres denote resolved and assigned amide N atoms that undergo $^{1}H/^{15}N$ chemical shift perturbations from 0 ppm (cyan) to 0.15 ppm (red), between the cell envelope and liposome environments. **(B)** Spheres denote resolved and assigned amide N atoms that undergo $^{1}H/^{15}N$ chemical shift perturbations from 0 ppm (cyan) to 0.20 ppm (blue), in the presence of NHS.

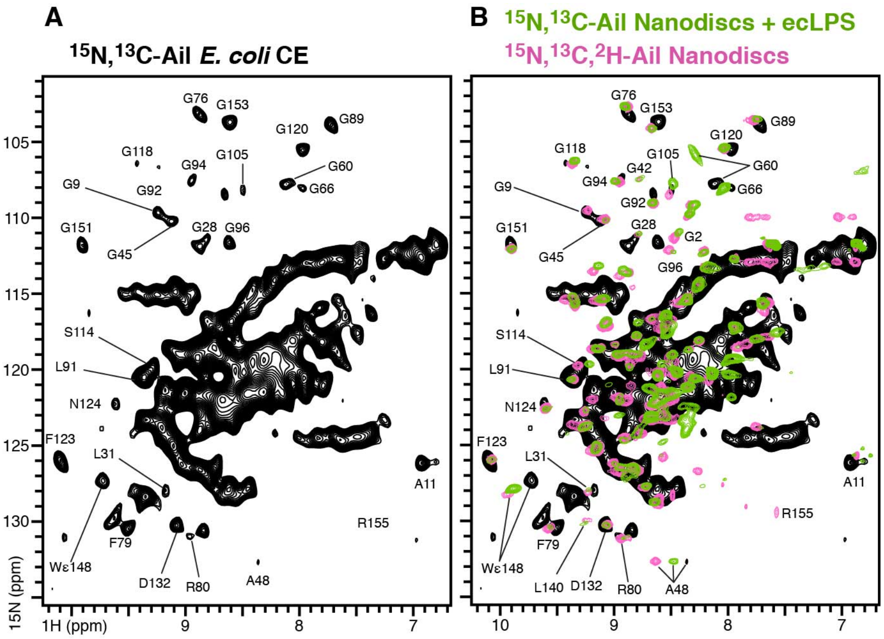


**Figure S5 Solid-state NMR $^1H/^{15}N$ CP-HSQC 2D spectra and solution NMR $^1H/^{15}N$ HSQC 2D spectra of Ail in *E. coli* cell envelopes (CE) and nanodiscs.** Solid-state NMR CP-HSQC spectra (black) were acquired for ($^{15}N$,$^{13}C$)-Ail in *E. coli* CE from Lemo21(DE3) cells, at 900 MHz, 30°C, with a MAS rate of 60 kHz and 1,600 transients. Solution NMR HSQC spectra were acquired for purified Ail uniformly labeled with $^{15}N$, $^{13}C$ and $^2H$ in lipid nanodiscs (pink), or uniformly labeled with $^{15}N$ and $^{13}C$ in nanodiscs reconstituted with *E. coli* rough-type LPS (green). The solution NMR spectra of Ail nanodiscs have been described previously [14, 19]

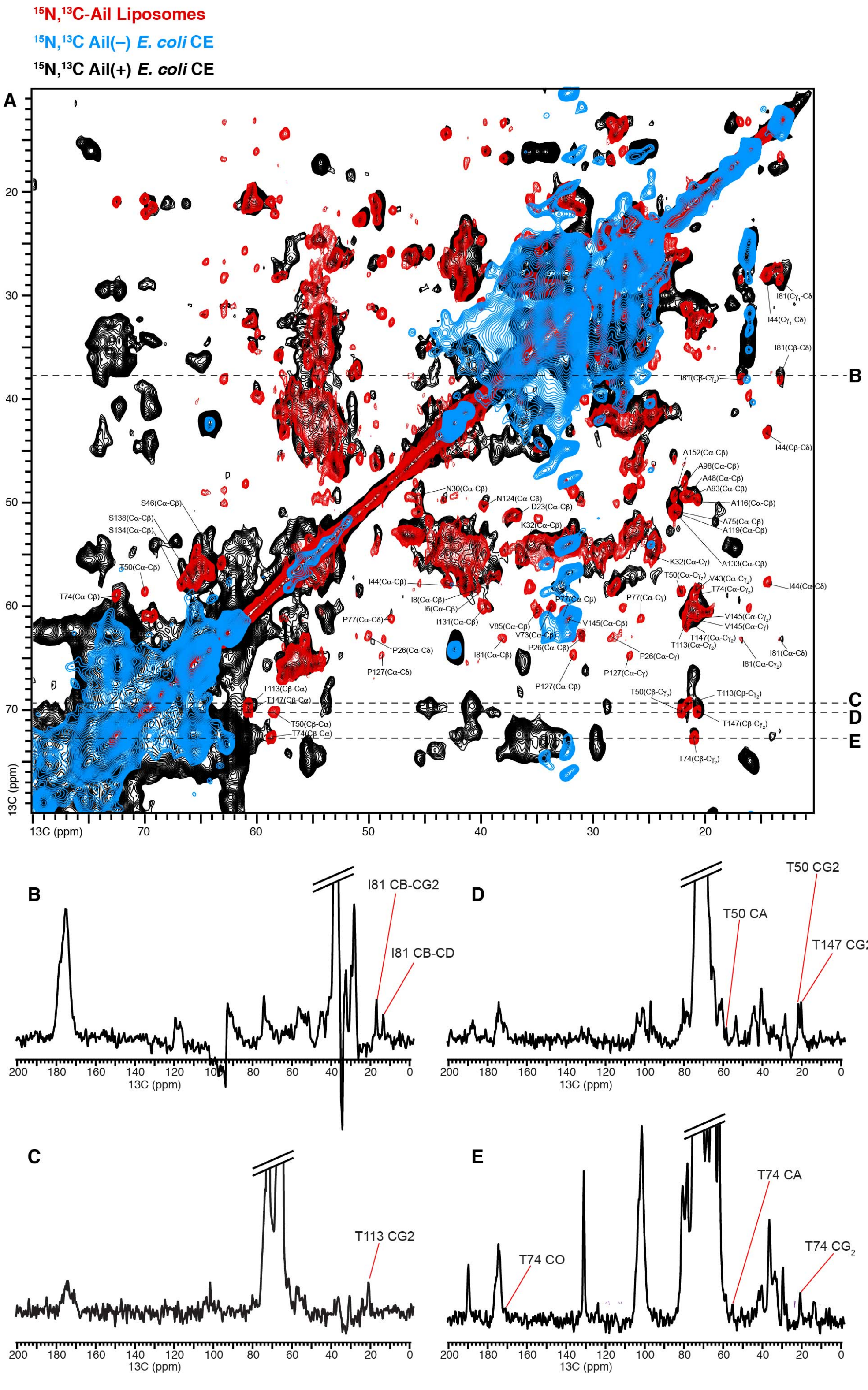


**Figure S6. $^{13}C/^{13}C$ PDSD solid-state NMR spectra. (A)** 2D Spectra were acquired at 750 MHz (204 t1 increments with 304 transients) for $^{15}N$,$^{13}C$-Ail in *E. coli* cell envelopes (black); 500 MHz (444 t1 increments and 256 transients) for $^{15}N$,$^{13}C$ Ail(–) *E. coli* cell envelopes (blue); and 750 MHz (512 t1 increments and 64 transients) for $^{15}N$,$^{13}C$-Ail in reconstituted liposomes (red). Resolved assigned Ail peaks are marked. The horizontal dashed lines mark the 1D spectral slices extracted from the spectrum. **(B-E)** 1D Slices taken from the 2D spectrum of Ail(+) cell envelopes..

**Table S1.** Resolved $^1H/^{15}N$ NMR signals with tentative assignments derived by spectral comparison with the solid-state and solution NMR spectra of purified Ail reconstituted in liposomes or nanodiscs (18, 19). Missing peaks are denoted by the letter "X". The combined difference (ΔHN) of amide $^1H$ and $^{15}N$ chemical shifts was calculated as: $\Delta HN = [(\Delta H)^2+(\Delta N/5)^2]^{1/2}$.

| Ail residue | liposomes | | CE | | CE + NHS | | lipo VS CE | CE ± NHS |
|---|---|---|---|---|---|---|---|---|
| | $^1H$ (ppm) | $^{15}N$ (ppm) | $^1H$ (ppm) | $^{15}N$ (ppm) | $^1H$ (ppm) | $^{15}N$ (ppm) | ΔHN (ppm) | ΔHN (ppm) |
| **9G** | 9.139 | 109.788 | 9.229 | 109.669 | 9.123 | 109.485 | 0.093 | 0.113 |
| **11A** | 6.849 | 126.575 | 6.931 | 126.083 | 6.847 | 126.337 | 0.128 | 0.098 |
| **23D** | 8.141 | 114.416 | 8.213 | 114.070 | 8.184 | 114.713 | 0.100 | 0.132 |
| **28G** | 8.736 | 111.021 | 8.809 | 111.373 | 8.731 | 111.717 | 0.101 | 0.104 |
| **45G** | 9.071 | 110.066 | 9.115 | 110.230 | 9.000 | 110.325 | 0.055 | 0.116 |
| **48A** | 8.324 | 132.972 | 8.358 | 132.585 | X | X | 0.085 | — |
| **60G** | X | X | 7.966 | 108.059 | 8.035 | 107.870 | — | 0.078 |
| **66G** | X | X | 8.110 | 107.786 | 8.186 | 107.972 | — | 0.085 |
| **76G** | 8.852 | 102.744 | 8.874 | 103.164 | 8.921 | 103.362 | 0.087 | 0.061 |
| **80R** | 8.810 | 131.135 | 8.833 | 130.502 | 8.800 | 130.396 | 0.129 | 0.039 |
| **81I** | 8.194 | 126.515 | X | x | X | X | — | — |
| **82N** | 7.837 | 112.360 | X | x | X | X | — | — |
| **89G** | 7.709 | 103.532 | 7.714 | 103.901 | 7.712 | 103.898 | 0.074 | 0.002 |
| **92G** | 8.567 | 108.949 | 8.649 | 108.458 | 8.564 | 108.557 | 0.128 | 0.088 |
| **94G** | 8.937 | 107.875 | 8.937 | 107.534 | X | X | 0.068 | — |
| **105G** | X | X | 8.487 | 108.189 | 8.487 | 107.165 | — | 0.205 |
| **118G** | 9.327 | 106.444 | 9.415 | 106.430 | 9.389 | 106.433 | 0.088 | 0.026 |
| **120G** | 7.969 | 105.521 | 7.967 | 105.562 | 7.945 | 105.220 | 0.008 | 0.072 |
| **123F** | 10.086 | 126.487 | 10.085 | 125.922 | 10.158 | 125.454 | 0.113 | 0.119 |
| **124N** | X | X | 9.604 | 122.208 | X | X | — | — |
| **131I** | 8.830 | 124.461 | 8.868 | 123.959 | 8.887 | 124.103 | 0.107 | 0.035 |
| **132D** | 9.013 | 130.121 | 9.068 | 130.170 | X | X | 0.056 | — |
| **151G** | 9.864 | 112.073 | 9.894 | 111.808 | 9.758 | 111.882 | 0.061 | 0.137 |
| **153G** | 8.630 | 103.914 | 8.602 | 103.716 | 8.641 | 103.971 | 0.048 | 0.064 |


18. Yao Y, Dutta SK, Park SH, Rai R, Fujimoto LM, Bobkov AA, Opella SJ, Marassi FM (2017) **High resolution solid-state NMR spectroscopy of the Yersinia pestis outer membrane protein Ail in lipid membranes**. *J Biomol NMR* 67, 179-190 (PMC5490241).

19. Dutta SK, Yao Y, Marassi FM (2017) **Structural Insights into the Yersinia pestis Outer Membrane Protein Ail in Lipid Bilayers**. *J Phys Chem B* 121, 7561-7570 (PMC5713880).
